# CASTOR: A machine learning platform for reproducible viral genome classification

**DOI:** 10.1101/082768

**Authors:** Mohamed Amine Remita, Ahmed Halioui, Abou Abdallah Malick Diouara, Bruno Daigle, Golrokh Kiani, Abdoulaye Baniré Diallo

## Abstract

**Motivation:** Advances in cloning and sequencing technology yielded a massive number of genome of virus strains. The classification and annotation of these genomes constitute important assets in the discovery of genomic variability, taxonomic characteristics and disease mechanisms. Existing classification methods are often designed for a well-studied virus. Thus, the viral comparative genomic studies could benefit from more generic, fast and accurate tools for classifying and typing newly sequenced strains of diverse virus families.

**Results:** Here, we introduce a fast, accurate and generic virus classification platform, CASTOR, based on a machine learning approach. CASTOR is inspired by a well-known technique in molecular biology: Restriction Fragment Length Polymorphism (RFLP). It simulates the restriction digestion of genomic material by different enzymes into fragments in-silico. It uses two metrics to construct feature vectors for machine learning algorithms in the classification step. We benchmark CASTOR for the classification of distinct datasets of Human Papillomaviruses (HPV), Hepatitis B Viruses (HBV) and Human Immunodeficiency viruses (HIV). Results reveal true positive rates of 99%, 99% and 98% for HPV Alpha species, HBV genotyping and HIV M group subtyping respectively. Furthermore, CASTOR shows a competitive performance compare to well-known HIV-specific classifier REGA and COMET on whole genome and *pol* fragments. With such prediction rates, genericity and robustness, as well as rapidity, such approach could constitute a reference in large-scale virus studies. Finally, we developed the CASTOR web platform for open access and reproducible viral machine learning classifiers.

**Availability:** http://castor.bioinfo.uqam.ca

## 1 Introduction

Genomic sequence classification assigns a given sequence into its related group of known sequences having similar properties, traits or characteristics. It is a fundamental practice in different research areas of microbiology yielding major challenges in comparative genomics. Accurate genomic sequence classification and typing help to have a better understanding of the evolution and phylogenetic relationships of viruses. They also help in determining pathogenicity, developing vaccines, studying epidemiology and drug resistances (Struck *et al.*, 2014). Recent advances in DNA sequencing and molecular biology techniques provide an immense collection of genomic information. Such data volume raises challenges for genetic-based classification techniques. Three main approaches have been designed and implemented to classify different types of viruses based on their genomic sequence characteristics. The first is *sequence alignment-based* approach which is widely used, e.g.: in similarity search methods (BLAST (Altschul *et al.*, 1997), USEARCH (Edgar, 2010), etc.) and in pairwise distance based-methods (PASC (Bao *et al.*, 2014), DEmARC (Lauber and Gorbalenya, 2012), etc.). The second is *phylogenetic-based* approach. It is implemented in several tools, e.g.: REGA (de Oliveira *et al.*, 2005) and Pplacer (Matsen *et al.*, 2010). The aim of these methods is to place an unknown sequence on a phylogenetic tree of a reference sequences. Each time a given sequence has to be classified, it is realigned with the set of reference sequences. Then, either a new phylogenetic tree is inferred or the given sequence is placed in the existing tree. The third is *alignment-free* approach including methods based on nucleotide correlations (Liu *et al.*, 2008) and sequence composition (Yu *et al.*, 2013; Struck *et al.*, 2014). It transforms sequences or their relationships to feature vectors and then constructs a phylogeny, statistical or machine learning model (Vinga and Almeida, 2003; Bonham-Carter *et al.*, 2013). These methods are reviewed in Vinga and Almeida (2003), Mantaci *et al.* (2008), Xing *et al.* (2010) and Bonham-Carter *et al.* (2013). Restriction fragment length polymorphism (RFLP), a molecular biology technique (Williams, 1989), is used to type different virus strains (Bernard *et al.*, 1994; Nobre *et al.*, 2008). Several computational and algorithmic approaches have tackled theoretical and experimental problems related to the restriction enzyme data such as phylogeny estimation (Felsenstein, 1992), SNP genotyping (Chang *et al.*, 2010) and analysis of RFLP digitized gel images (Maramis *et al.*, 2011). However, large-scale computational sequence classification based on the RFLP technique is not yet covered in literature. Due to the genetic polymorphism in DNA sequences, fragments resulting from enzyme digestions are different in terms of number and length between individuals or types. A set of restriction enzymes grounds a fragment pattern signature for each sequence. Therefore, similar sequences ought to have similar fragment patterns and thus similar restriction site distributions. This *a priori* knowledge could be used to build a machine learning model where sequences are represented by restriction site distributions as a feature vector and a class feature corresponding to a taxonomic level (genus, species, etc.). In this paper we introduce CASTOR, a machine learning web platform, to classify and type sequences. CASTOR integrates a new alignment-free method based on the RFLP principle. Our *in silico* method is independent of the sequence structure or function and is also not organism-specific. CASTOR is designed to facilitate the reuse, sharing and reproducibility of sequence classification experiments.

## 2 Material and methods

### 2.1 Overview of the approach

In this study, we propose an *in silico* approach to identify and classify viral DNA sequences based on their restriction enzyme sites using supervised machine learning techniques. Like other supervised learning approaches, ours is divided into two main units (Fig. 1). The *classifier construction unit* builds and trains classification models (or classifiers). It requires a set of reference viral genomic sequences, their classes and a list of restriction enzyme patterns. It starts by creating a training set including an ensemble of feature vectors. The latter is computed from the distribution of the restriction site patterns on the given DNA sequences and then refined by feature selection methods. A collection of learning classifiers are then trained and evaluated using 10-fold cross analyses in order to choose the best classifier. The second unit (*prediction unit*) is intended to predict classes or annotions of given viral sequences. The input data of this unit are a classifier, a set of DNA sequences and the same list of restriction enzyme patterns used to train the classifier.

**Fig. 1.**
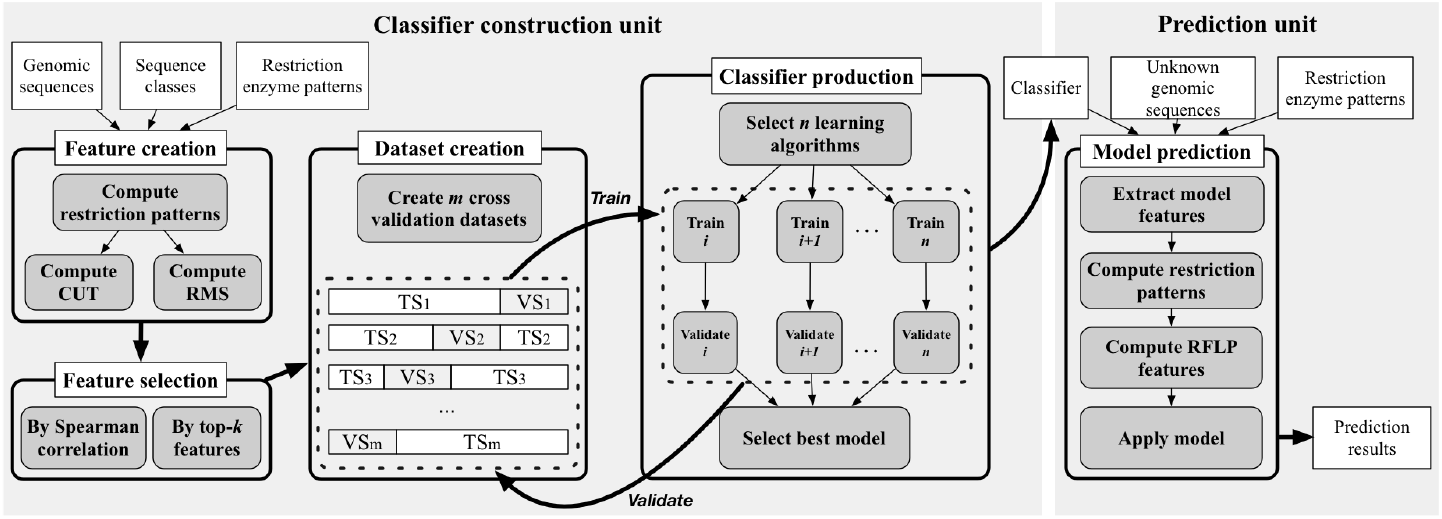
Overview of CASTOR kernel architecture. The kernel is composed of two main units (model construction and prediction). White rectangles represent input and output data; grey and curved rectangles represent processes. TS and VS are respectively training set and validation set.

### 2.2 Restriction fragment pattern-based features

In this study, we propose a set of features simulating the outcome of the RFLP technique. From REBASE database (Roberts *et al.*, 2015), we extracted a list of 172 type II restriction enzymes and their recognition sites. Type II family cleaves (cuts) DNA sequences precisely on each occurrence of the recognition site. Then, the restriction digestion of DNA sequences is computationally simulated. In order to build a training set, for each sequence *s* and enzyme *z* we compute two metrics representing the distribution of the digested fragments: the number of cuts of the enzyme (*CUT*(*s, z*)) and the root mean square of digested fragment lengths (*RMS*(*s, z*)) calculated as 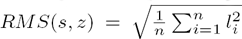 where *n* is the number of fragments (*CUT*(*s, z*) + 1) and *l*_*i*_ is the length of the *i*^*th*^ fragment in linear genomes. For circular genomes *n = CUT*(*s, z*). Other metrics could be easily computed from the fragment digestion to construct the feature vectors.

### 2.3 Feature selection methods

Selection of an optimal subset of features improves the learning efficiency and increases the predictive performance. Feature selection techniques reduce the learning set dimension by pruning irrelevant and redundant features. Two relevant methods of feature reduction are provided. The first method (*topAttributes*) ranks the features according to their information gain (Ben-Bassat, 1982) and a subset of top-*k* features is selected. Information gain estimates the mutual information between a feature and the target class. The second method (*correlation*) uses the Spearman’s rank correlation coefficient to construct a set of uncorrelated features. The *correlation* coefficient between two feature ranking vectors *u* and *v* of size *n* is computed as follows: 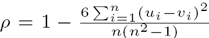. A two-tailed *p-value* is computed to test the null hypothesis which states that two feature vectors are uncorrelated. In order to compare and remove one of two correlated features, two methods could be used: feature with the largest sum of absolute correlation coefficients, or feature with the smallest information gain ranking.

### 2.4 Learning and evaluation

We explored three types of classifiers: (1) symbolic; using a C4.5 decision tree (J48) (Quinlan, 1993) and random forests (RFT) (Breiman, 2001), (2) statistical; using a naive Bayes classifier (NBA) (Langley *et al.*, 1992; John and Langley, 1995), a support vector machine (SVM) (Cortes and Vapnik, 1995) and K-nearest neighbors (IBK) (Cover and Hart, 1967; Aha *et al.*, 1991) and (3) Meta-learners; using Adaboost (ADA)(Freund and Schapire, 1997) and Bagging (BAG) (Breiman, 1996) both combined with J48 (see Table S1). A 10-fold cross-validation strategy is used to assess the performance of the trained classifiers. For each class, a set of performance measures is computed and averaged from all folds. Performance measures are weighted according to the number of instances and computed for the overall classification. The performance measures are: *TPR* = *TP*/(*TP* + *FN*), *FPR* = *FP*/(*FP* + *TN*), *Precision* = *TP*/(*TP* + *FP*) and *F – measure* = 2 × *TPR* × *Precision*/(*TPR* + *Precision*) where *TP, TN, FP,* and *FN* are the number of true positive, true negative, false positive and false negative predictions, respectively. *TPR* and *FPR* are true positive rate and false positive rate, respectively.

### 2.5 Datasets

We applied our approach to study the classification of three distinct viruses: Human PapillomaVirus (HPV), Hepatitis B Virus (HBV) and Human Immunodeficiency Virus type 1 (HIV-1). 1) HPVs have a circular double stranded DNA genome of ~8000bp belonging to five genera (Alpha, Beta, Gamma, Mu and Nu). HPVs belonging to a genus share over 53% identity of their complete genomes (CGs) and HPVs in the same species level share over 62% identity (Daigle *et al.*, 2015; Bernard *et al.*, 2010). We assess the approach performance for the classification of HPVs in the genus and species taxonomic levels. At the species level, we selected only the Alpha HPV genus representing the more abundant and the most diverse genomes in databases. It is divided into thirteen species (Alpha 1-11, Alpha 13-14). Unfortunately some HPV genera (Mu and Nu) and Alpha HPV species (1, 5, 8, 11 and 13) were underrepresented and were discarded. 2) HBV genomes are smaller (3200bp) and are circular partly double stranded DNA. HBVs are classified into eight genotypes (A-H) with at least 8% divergence between their genomic sequences (Schaefer, 2007). We evaluated the CASTOR performances for the genotyping of HBV strains. HPV and HBV genomic sequences were downloaded from NCBI RefSeq database (Coordinators, 2016). Only complete, clean and well-annotated sequences were selected. The taxonomic annotations were extracted from NCBI Taxonomy database (Coordinators, 2016). 3) HIV-1 genome has two copies of positive-sense single-stranded RNA with ~9700bp for each. Phylogenetically, HIV-1 strains are divided into four groups: M, N, O and P (Robertson *et al.*, 2000; Plantier *et al.*, 2009). M group strains are worldwide prevalent. They are categorized into pure subtypes (A-D, F-H, J and K) and recombinant forms (up to 70 CRFs and URFs). Genetic variations between subtypes are about 20-30% for *env* gene, 7-20% for *gag* gene and 10% for *pol* gene (Gao *et al.*, 1998). For HIV-1 classification, we studied CGs and fragments covering *pol* gene from the position 2253 to 3554 with respect to HXB2 reference sequence and having a minimum size of 1Kbp. HIV-1 sequences were extracted from the Los Alamos HIV database (http://www.hiv.lanl.gov/). Each class ought to have an adequate number of genomic sequences in order to have a representative genetic diversity.

### 2.6 Simulation studies

In order to identify the best parameters for tuning the classifiers, we randomly divided into 10 sets each, the HPV genera, HPV Alpha species, HBV genotypes datasets. For each obtained datasets, we performed a 10-fold cross-validation studies with different classifiers constructed as follows. We constructed all the combinations of the two metrics (*CUT* and *RMS*), the two sets of feature selection techniques (including *topAttributes* with *top – k* = 10, 50, 100, 172; *correlation* with *ρ* = 0.5, 0.7, 1,*p – value* = 0.05, 0.005, 0.5*E* – 5 thresholds and two methods to eliminate correlated features) resulting to 22 combinations and seven learning algorithms. This construction yielded 308 *combinations \** 10 *datasets* = 3080 *experiments* for each virus classification (see Figures S1 and S2). With the best set of parameters in the feature selection models (*topAttributes: top – k* = 100 and *correlation* parameters: *ρ* = 1, *p – value* = 0.5*E* – 5 and information gain as elimination method), we performed a second simulation study for the HPV genera, HPV Alpha species, HBV genotype, HIV-1 M subtypes (CGs) and HIV-1 M subtypes (*pol* fragments). Hence, in this simulation, we drawn the combination of 2*metrics\**2 *feature selection methods\**7 *learning algorithms\** 10 *datasets* = 280 *experiments* for each virus classification. This constitutes the main experiments presented in the result section. Raw viral sequence datasets constructed above were class-size imbalanced, i.e., the difference in the number of genome sequences belonging to each class was relatively large. Under-sampling (down-sizing) majority class approach has been shown to perform well (Blagus and Lusa, 2010) and could be used with standard algorithms. Hence, from each previous dataset, we randomly performed under-sampling of the larger classes and without replacement to have relatively the same size of the other classes. The interval of sampling size is given in each result tables.

## 3 Results and discussion

### 3.1 Classification with RFLP signatures in virus families

Figure 2 highlights the natural RFLP cuts in the collected HPV, HBV and HIV-1 datasets. The second column of the figure shows the multidimensional scaling (MDS) plot of the first two dimensions of the distances between the feature vectors of the genomes. The separation between the different HPV genera (Fig. 2a) could approximatively be drawn, which is partly the case for the HPV species. The *Cohesion* (Daigle *et al.*, 2015) and *Silhouette* (Rousseeuw, 1987) indices allow to measure the compactness and separability of classes. Here, both indexes show moderate values (between 0.2 and 0.8 for *Cohesion index* and −0.2 to 0.7 for *Silhouette index*) indicating that the classes are not really crisp. Several instances could be either mis-labeled or share the same RFLP cut patterns with other classes resulting in low or negative values of *Silhouette index* in HPV Alpha 3, 7 and HPV Gamma (Fig. 2a). With CASTOR, the best HPV Alpha Species classification obtains a *TPR* of 0.992 and *FPR* of 0.002 in 10-fold cross validation analyses of 118 instances (see Table 1). The power of RFLP cuts in classification of viruses could be observed in HBV genotypes heatmap (see Fig. 2b). HBV highlights three genotypes (A, E and F) with *Cohesion indexes* for most instances above 0.7 indicating very coherent classes. The *Silhouette index* plots show several instances of B, C, E and G genotypes that have an important disagreement with their affected classes (*Silhouette index* < −0.1). Even with these constraints, CASTOR achieves the genotyping of 230 HBV instances with *TPR* of 0.996 and *FPR* of 0.001 according to a 10-fold cross validation study (see Table 1). The HIV-1 cut site patterns have more variability among pure subtypes and CRFs (Fig. 2c). This variability among classes is reflected on the low values of the *Cohesion index* (<= 0.4) All, suggesting either variability, noise or mislabels. For instance, > 30% of HIV-1 B and HIV-1 C instances tend to have RFLP cut pattern of another subtypes (negative *Silhouette indexes*). With CASTOR, the subtyping of HIV-1 group M within 18 main subtypes was assessed for 597 instances with a *TPR* of 0.983 and *FPR* of 0.001. Previously, it has been clearly shown that RFLP has a power for classification in several viruses (Bernard *et al.*, 1994; Nobre *et al.*, 2008). But these studies are mostly limited to two to five classes. To the best of our knowledge, our study constitutes the first large scale and multi-class analyses of RFLP cut for classification.

**Fig. 2.**
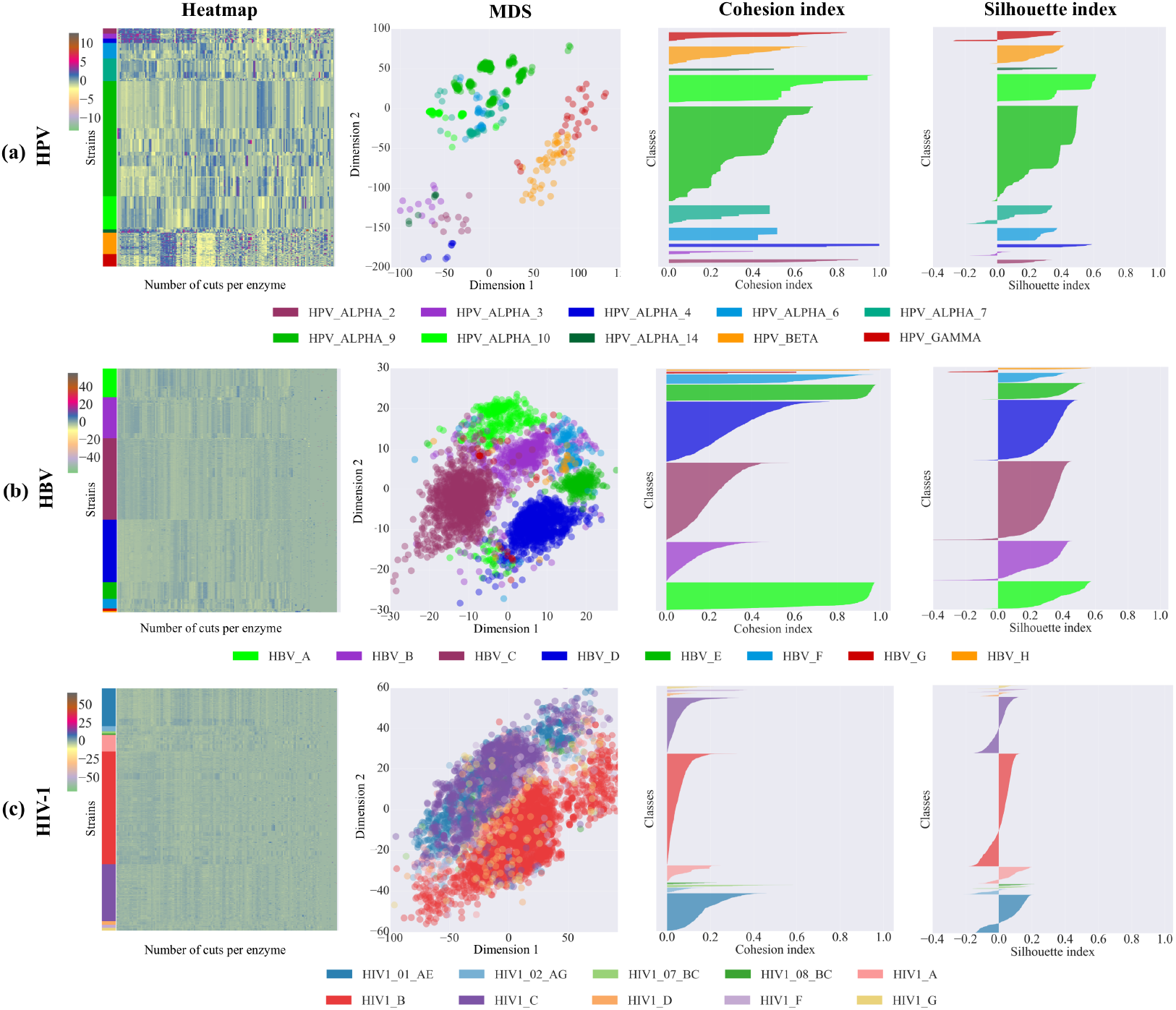
Class cohesion of three virus datasets. The four columns illustrate the separability and compactness of three virus complete genomes datasets based on 172 restriction enzyme RFLP cuts. The first column shows heatmaps of CUT clustered by x-axis. The samples in the y-axis are grouped by studied classes followed by intra-class clusterings. The second column shows MDS of the CUT distances between samples. The third and fourth column represents, respectively, the Cohesion and Silhouette indices of the classes. (a) Classes in HPV are Alpha species, Beta and Gamma genera. (b) Classes in HBV are A-H genotypes (c) Classes in HIV-1 are M pure subtypes and CRFs. The first 10 largest classes for each dataset (except HBV).

**Table 1.** CASTOR best accuracies on the classification of five datasets.

| Type of virus | Organism | Classification | # of classes | # of instances | *TPR* | *FPR* | *F-measure* | Classifier ID |
| --- | --- | --- | --- | --- | --- | --- | --- | --- |
| Group I dsDNA | HPV | Genera | 3 | 125 | 0.992 | 0.005 | 0.992 | PMSHPV01 |
|  |  | Alpha species | 8 | 118 | 0.992 | 0.002 | 0.992 | PMSHPV02 |
| Group VII dsDNA-RT | HBV | Genotypes | 8 | 230 | 0.996 | 0.001 | 0.996 | PMSHBV01 |
| Group VI ssRNA-RT | HIV-1 | Groups | 4 | 76 | 1.000 | 0.000 | 1.000 | PMSHIV01 |
|  |  | M Subtypes | 18 | 597 | 0.983 | 0.001 | 0.983 | PMSHIV02 |

This table contains the best results of the experimental study performed on the different datasets. The evaluation measures are obtained with 10-fold cross validation analysis. The column Classifier ID contains the corresponding models available in CASTOR platform.

### 3.2 Machine learning classifiers tuning and performance

The CASTOR platform relies on machine learning methods for the classification of viruses based on RFLP signatures in nucleotide sequences. The platform is detailed in the CASTOR platform section. Three important parameters constitute the kernel of each CASTOR classifier (a metric, a feature selection method, a learning algorithm). To assess the different combination of the models, we performed a 10-fold cross-validation of the 280 experiments associated to each of the five datasets. From the overall results of the three virus datasets, it is tricky to distinguish the best candidate between *CUT* and *RMS* metrics. In the genotyping of HBV, *CUT* performs better than *RMS* (p-value = 0.0012) while in the HPV genera and species classifications *RMS* performs better than *CUT* (p-values 5.00E-03 and 0.0293 respectively) (Fig. S3). However the weighted mean *F-measures* for both methods are in all case >= 0.90 (with minimum of 0.79 and maximum of 0.99). The same analyses were performed on HIV-1 CGs and *pol* fragments. *CUT* and *RMS* perform quite similar in both datasets when comparing the mean weighted *F-measure* (non-significant p-values). Due to the variability of HIV-1, the mean weighted *F-measure* is 0.86 in CGs and 0.80 in *pol* fragments. Hence for the remaining of our study, we will fix the metric according to its performance on the corresponding datasets. Figure S4 presents the comparative analyses of the two feature selection methods in the 280 experiments for each dataset. The Wilcoxon/kruskal-wallis tests comparison of the mean of weighted *F-measure* for the two feature selection approaches show that *correlation* and *topAttribute* results are not statistically different in all datasets. In fact, the two methods are correlated in the three viruses with a Spearman’s rank correlation coefficient ranging between 0.77 and 0.96 (see Fig. S6). In these simulations, the seven learning algorithms have various performance according to the different datasets. The algorithm J48 has the worst weighted *F-measure* values (see Fig. 3). However, its performance improves when combined with RFT or BAG algorithms. In general, SVM performs better in 4/5 datasets with weighted mean *F-measure* > 0.95 and ranking number 1 in HPV Alpha species, HBV genotypes and HIV-1 subtypes classification and 4 in HPV genera classification. It is followed by RFT, NBA and IBK. However, RFT and NBA are affected by a large variance (Fig. 3). These ranking are more less observable on Figure S5 and S6. While most algorithms have similar performance with *CUT* or *RMS*, Naive Bayes surprisingly performs better with *CUT*.

**Fig. 3.**
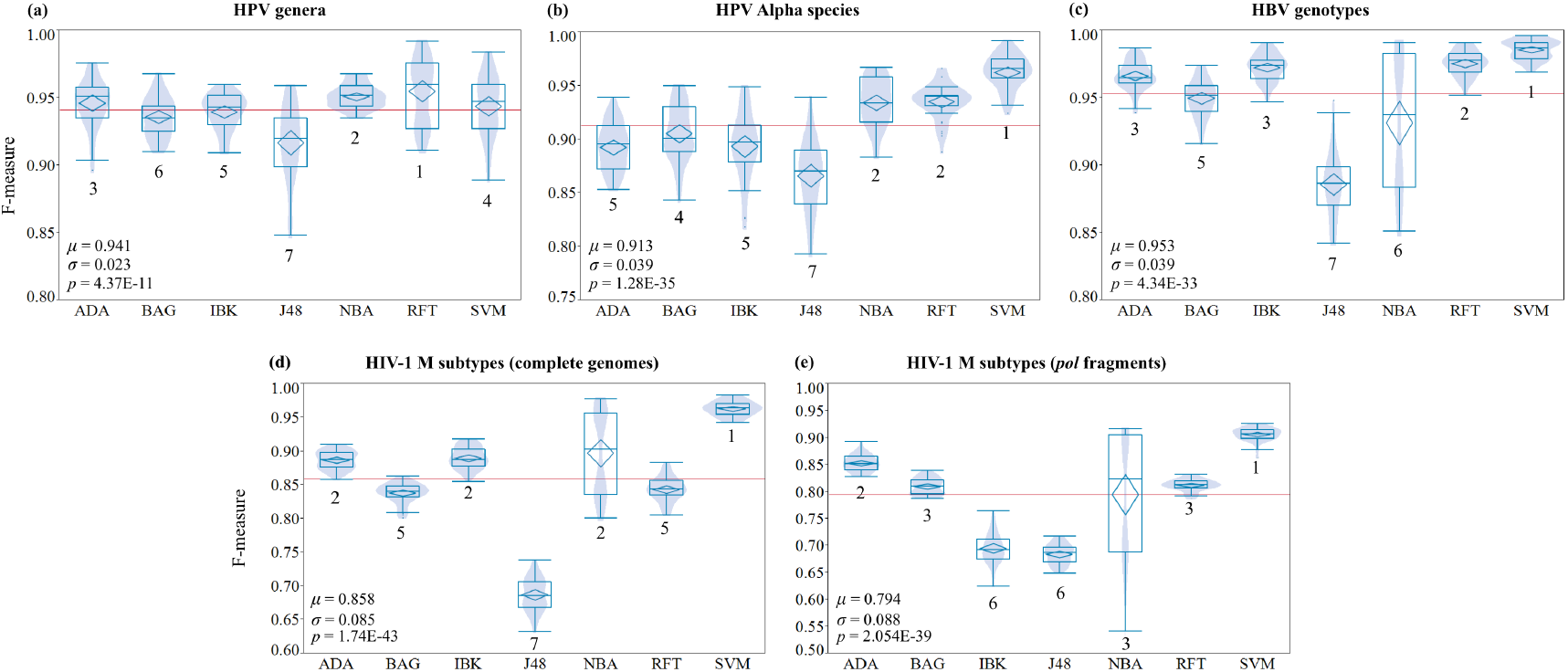
Learning algorithm evaluation on five datasets. This figure illustrates the F-measure distribution (boxplot) of seven learning algorithms on the prediction for (a) HPV genera, (b) HPV Alpha species, (c) HBV genotypes, (d) HIV-1 M subtypes with complete genomes (e) HIV-1 M subtypes with pol fragments. HPV and HBV datasets are complete genomes. The number below each boxplot corresponds to the statistically discriminative ranking of the algorithms. The ranking is performed with paired Student’s t test. *µ*, *σ* are the mean and the standard deviation of the overall F-measures. p is the p-value of the statistically significance of the F-measure median differences among the algorithms computed with the Wilcoxon signed rank test.

### 3.3 Assessing the performance CASTOR on HIV data

Table 2 highlights Castor prediction accuracies on five CG and seven *pol* sequence fragments based HIV-1 classification. The *TPR* of the best classifier for the main HIV-1 types M, N, O and P indicates that all the sequences are correctly classified. For the prediction between the main HIV-1 pure subtypes as well as CRFs, are above 0.98 (with *FPR* <= 0.001) for both CGs and *pol* fragments when the pure subtypes and CRFs are separate models. When combining Pure subtypes and CRFs, the *TPR* still remains above 0.98 for CGs but it drops at 0.92 when the classes are balanced to 30 instances per class or 0.96 for 200 instances per class. It appears that the CASTOR models are underperforming when we try to predict between pure subtypes and CRFs (*F-measure* of 0.795 and 0.885 for CGs and *pol* fragments respectively).

**Table 2.** Evaluation of HIV classification with CASTOR

|  | Classification | # of classes | # of instances | [min - max] instances/class | TPR | FPR | F – measure | Classifier ID |
| --- | --- | --- | --- | --- | --- | --- | --- | --- |
| Complete genomes | Groups (M, N, O and P) | 4 | 76 | [4 - 32] | 1.000 | 0.000 | 1.000 | PMVHIVGC01 |
|  | Pure subtypes | 6 | 189 | [30 - 36] | 0.995 | 0.001 | 0.995 | PMVHIVGC02 |
|  | CRFs | 12 | 234 | [10 - 30] | 1.000 | 0.000 | 1.000 | PMVHIVGC03 |
|  | Pure subtypes and CRFs | 18 | 423 | [10 - 36] | 0.981 | 0.001 | 0.981 | PMVHIVGC04 |
|  | Pure subtypes vs CRFs | 2 | 200 | [100 - 100] | 0.795 | 0.205 | 0.795 | PMVHIVGC05 |
| *pol* fragments | Groups (M, N, O and P) | 4 | 94 | [4 - 45] | 1.000 | 0.000 | 1.000 | PMVHIVPL01 |
|  | Pure | 6 | 1800 | [300 - 300] | 0.983 | 0.003 | 0.983 | PMVHIVPL02 |
|  | CRFs | 16 | 480 | [30 - 30] | 0.971 | 0.002 | 0.971 | PMVHIVPL03 |
|  | CRFs | 6 | 1200 | [200 - 200] | 0.993 | 0.001 | 0.993 | PMVHIVPL04 |
|  | Pure subtypes and CRFs | 23 | 690 | [30 - 30] | 0.920 | 0.004 | 0.919 | PMVHIVPL05 |
|  | Pure subtypes and CRFs | 12 | 2400 | [200 - 200] | 0.962 | 0.003 | 0.962 | PMVHIVPL06 |
|  | Pure subtypes vs CRFs | 2 | 200 | [100 - 100] | 0.885 | 0.115 | 0.885 | PMVHIVPL07 |

This table contains the TPR, FPR and F-measure of 12 HIV classifications obtained with 10-fold cross validation analysis. For each classification, the number of corresponding classes and instances are given. The range [min-max] indicates the interval of instance frequencies per class used during the training of each model. The column Classifier ID contains the corresponding models available in CASTOR platform.

Next, we also compare the performance of CASTOR against the most powerful and widely used specific HIV-1 predictors namely COMET (Struck *et al.*, 2014) and REGA vs2 (de Oliveira *et al.*, 2005) (Figure 4). It is important to notice that these programs are fixed and do not allow neither any changing on the trained classes nor new training samples. Here the actual training of COMET and REGA includes respectively 55 and 22 classes. To avoid under-represented classes, CASTOR was trained on 18 classes for CGs and 28 classes for *pol* fragments (models are available under the classifier idPMSHIV02 and PMSHIV03, respectively). We performed three comparisons (*complete sampling, specific subtypes, common subtypes*; see Figure 4). REGA performs the best for CGs when COMET outperforms for *pol* fragments. But their performance drastically dropped in the other analyses by more than 10% compared to the best performing method and arriving at the third position (Figure 4). Meanwhile CASTOR is second in both two datasets. In CGs, CASTOR obtained a correct classification of 72.41% against the sampling of LANL data when REGA obtains 76.77%. But when testing predictors on their trained classes, the percentage of correct classification drastically increases to 98.33% and 96.61% respectively for REGA and CASTOR. This result remains almost the same when comparing only the common trained class among the three predictors (Figure 4). These common classes cover 75% and 93% of the overall instances of the sampling of CGs and *pol* fragments, respectively. CG data includes 6 classes with 4 pure subtypes and 2 CRFs (Table S2). The mean *TPR* of CASTOR is higher than 0.95 in either pure subtypes or CRFs. The *TPR* of REGA drops to 0.83 when assessing CRFs and remains almost perfect for pure classes (Table S2).

**Fig. 4.**
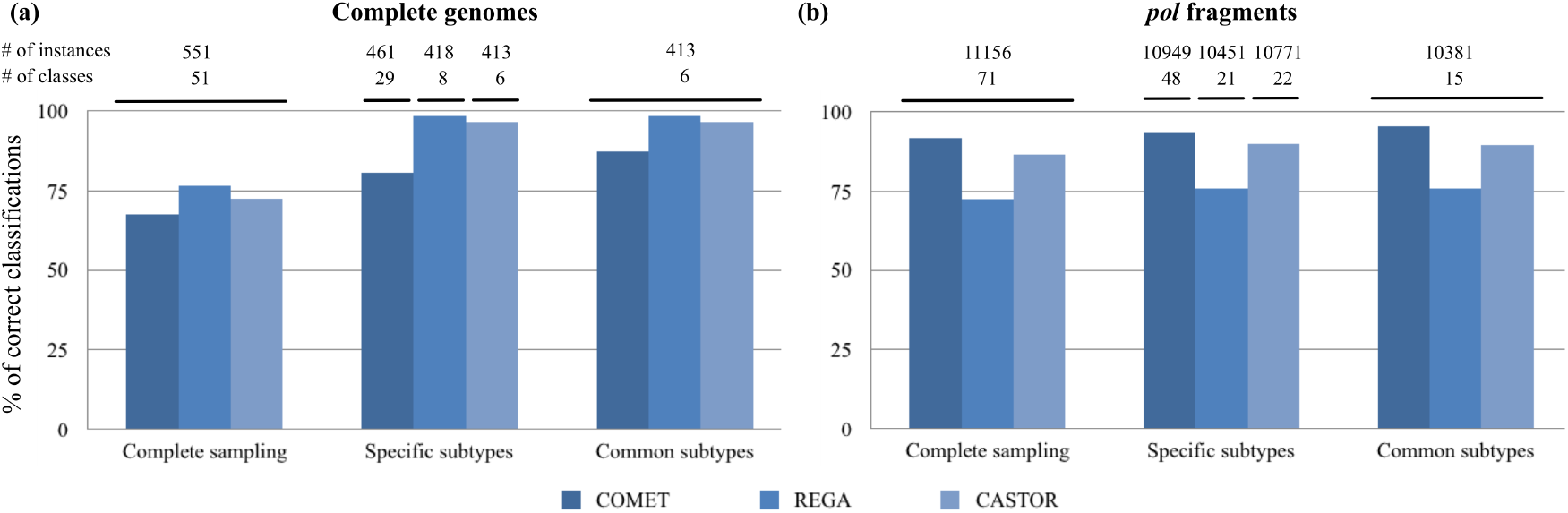
Performance of CASTOR with COMET and REGA predictors on HIV-1 datasets. The panels (a), (b) show the percentage of correct classifications for HIV-1 complete genomes and HIV-1 pol fragments, respectively. The number of instances and the associated classes for each sampling is presented above the panels. Complete sampling corresponds to 10% of LOS Alamos HIV-1 data selected randomly. In specific subtypes sampling, the predictors are assessed against their trained classes. In common subtypes sampling, the predictors are assessed against the classes intersection between the three trained predictors.

In *pol* fragments, COMET outperforms CASTOR and REGA in all comparisons. Against the 10% random sampling of LANL, COMET, REGA and CASTOR have respectively a percentage of correct classification of 91.74%, 72.48% and 86.64%. This picture is confirmed when comparing only the common trained classes where COMET reaches 95.57% and CASTOR 89.51%. Notice that REGA could not perform higher than 76% and has a mean *TPR* of 0.96 in pure subtypes competing with COMET. In CRF instances, COMET and CASTOR obtain an equal mean of *TPR* at 0.81 (Table S3). However, CASTOR has higher *FPR* that is reflected on the mean *F-measure* of 0.77 compare to 0.84 for REGA. The fact that *FPR* values are higher in CASTOR compare to the two other programs are not surprising. Since REGA and COMET are specifically tuned to predict HIV data, their predictions with lower scores tend to be discarded or ambiguous. For instance COMET has 32% of its CG prediction that is unassigned as well as 5% of its *pol* fragment predictions. Hence, these numbers are higher than the false positive values of CASTOR, but there are not included in the *FPR* computation. But, it will be interesting to include in CASTOR a threshold of inclusion of a given sequence into a class. This could help reducing the *FPR* but it would necessitate deeper analyses. It should also be associated to the *open-set* classification problem that is beyond the scope of this paper. Even though CASTOR is not a specific HIV-1 classifier, it competes with the most powerful method in HIV-1. Unlike COMET and REGA, CASTOR provides an easy way of performing several types of classification (see Table 2). It also has no restriction in the size of data and is really time efficient. Hence, we completed the analysis by performing a test on whole LANL. For CGs (3 778 instances), CASTOR computes the test in 1m59s with and accuracy of 91.2%. While for the *pol* fragments (119 005 instances), it requires 20min10sec with an accuracy of 85.41%. It shows that CASTOR takes 0.01sec to process a sequence that is far more efficient than the time results indicated in (Struck *et al.*, 2014) for REGA (28sec/sequence), but 10-fold less efficient than COMET (0.001sec/sequence) (Struck *et al.*, 2014). Furthermore, due to size issues, it is not possible to perform such large analyses in actual version of COMET server. Overall, CASTOR highlights good accuracy on the classification of the three studied viruses. However this accuracy is slightly lower than specific virus predictors as shown previously. But it exhibits more analysis capacity, permitting several and highly accurate set of classifications. As shown in 2, this accuracy is higher than 90% for almost all studies except for comparing HIV-1 Mpure subtypes vs CRFs. For less complex genomes such as HPV and HBV, the weighted mean *F-measure* is higher than 0.96. It will allow to increase the class representatives, to add or remove classes and also to benchmark several types of classification. For viruses that no specific classifier exists, it could accurately cover the needs as it is for HPV, instead of relying on the closes sequence search such as BLAST (Altschul *et al.*, 1997) or USEARCH (Edgar, 2010). Sequence search is generally not recommended for subtyping since it will not allow the identification of novel forms, it cannot also aggregate common attributes of a class while predicting (Struck *et al.*, 2014; Edgar, 2010).

### 3.4 CASTOR web platform

CASTOR is available as a public web platform. It is composed of four main applications. (1) **CASTOR-build** allows a user to the create and train new classifiers from a set of labeled virus sequences. It contains default parameters and advanced options letting a user to customize the classifier parameters. It can be used also to update the parameters or input sequences of an already built classifier. The constructed classifiers can be saved in an exportable file locally or publish to the community via CASTOR-database described below. (2) **CASTOR-optimize** constructs improved classifiers. unlike CASTOR-build that allows user to define metrics, algorithms and feature selection models, It assesses all combinations of the classification parameters and provides the best fitting classifier according to the input data. (3) **CASTOR-predict** is the kernel application that allows user to annotate a viral sequences according to a chosen classifier. It also serves as evaluation module for classifiers with a labeled test sets. The results are provided with enriched graphics and performance measures (4) **CASTOR-database** is a public database of classifiers which allow the community to share their expertise and models. It facilitates experiment reproducibility and models refinement. Asearch engine and classifier properties viewer are also implemented. Hence, from the interface of CASTOR-database, users can download, reuse, update and comment the published classifiers. In the best of our knowledge, this platform constitutes the first RFLP prediction based platforms for the classification of viral sequences.

## 4 Conclusion

In this paper, we have shown that RFLP has a great performance in large scale sequence classification such as typing, subtyping, genotyping and others. We also provide CASTOR, the first generic viral sequence classification platform based on RFLP. We raised that CASTOR can perform well in different type of viruses Group I, Group VI and Group VII (see Table 1) with weighted mean *F – measure* > 0.90 in most of the case. In the future, we will attempt to increase the performance by modelling the boundaries of the classes and including *open-set* approach to deal with instances from unknown classes. The CASTOR platform implements several metrics and classifiers, allowing generic and diverse analyses within the same environment. CASTOR allows the storage of models allowing for reproducible experiments and open data access. Even though, CASTOR is scale for viruses, it can be used and extend easily for other type of organisms, including whole genome and partial sequences. In the future, more models will be included, in particular those for less studied organisms and/or without dedicated tools. Moreover, scientists could add their tuned models helping CASTOR to enhance the predictions. We will also optimize the platform to allow diverse type of classification such as functional, disease related, geographical classifications. Hence, CASTOR could quickly become a reference in comparative genomics focusing on various type of sequence classification.

